# Functional specialization of specific DC subsets in the synovial fluid of JIA patients

**DOI:** 10.1101/2022.02.09.479707

**Authors:** Arjan Boltjes, Anoushka Ashok Kumar Samat, Maud Plantinga, Michal Mokry, Bas Castelijns, Joost F. Swart, Bas J. Vastert, Menno Creyghton, Stefan Nierkens, Jorg van Loosdregt, Femke van Wijk

**Author notes:** Equal contribution (Arjan Boltjes, Anoushka Ashok Kumar Samat, Maud Plantinga). Correspondence: Dr. F. van Wijk, KC02.085.2, Center for Translational Immunology, University Medical Center Utrecht, P.O. Box 85090, 3508 AB Utrecht, the Netherlands.

## Abstract

Dendritic cells (DC) are crucial for initiating and shaping immune responses. So far, little is known about heterogeneity and functional specialization of human DC subsets in (local) inflammatory conditions. We profiled conventional (c)DC1, cDC2 and monocytes based on phenotype, transcriptome and function from a local inflammatory site, namely synovial fluid (SF) from patients suffering from a chronic inflammatory condition, Juvenile Idiopathic Arthritis (JIA). cDC1, a relatively small dendritic cell subset in blood, are strongly enriched in SF. SF cDC1 showed a quiescent immune signature without a clear inflammatory profile low expression of PRR, chemokine, and cytokine receptors, and poor induction of T cell proliferation and cytokine production, but selective production of IFNλ upon polyinosinic:polycytidylic acid exposure. In stark contrast, cDC2 and monocytes from the same environment, showed a pro-inflammatory transcriptional profile, high levels of (spontaneous) pro-inflammatory cytokine production, and strong induction of T cell proliferation and cytokine production, including IL-17. Although the cDC2 and monocytes showed an overlapping transcriptional core profile, there were clear differences in the transcriptional landscape and functional features, indicating that these cell types retain their lineage identity in chronic inflammatory conditions. In conclusion, our findings suggest that at the site of inflammation, there is specific functional programming of human DC subsets, especially cDC2. In contrast, the enriched subset of cDC1 remains relatively quiescent and seemingly unchanged under inflammatory conditions, pointing to a more regulatory role.

## Introduction

Dendritic cells (DC) play a pivotal role in the induction and maintenance of the immune response, linking innate and adaptive immunity^1^. Together with monocytes, they form a heterogeneous group of professional antigen-presenting cells (APC) cells that display characteristic functional properties that together regulate the immune system towards both immunity or tolerance^2,3^. Although the maintenance of DC subsets is undergoing consistent dynamic change, monocytes, conventional DC (cDC) and plasmacytoid DC (pDC) are widely accepted as the most dominant APCs in steady state peripheral blood (PB)^4–6^. The PB DC subsets are defined as CD141^+^cDC1, CD1c^+^cDC2 and CD123^+^ pDCs^7^. DCs are also present in tissues where they display a more mature phenotype than their blood counterparts^8,9^. However, despite this maturation difference, the main DC subsets align across tissues and species in steady state^5,10,11–13^. Within these populations there is additional heterogeneity depending on, amongst others, developmental stages^10^. In inflammatory settings APC phenotype and function can change^8^. It remains poorly understood how the APC subtypes are functionally programmed at the site of inflammation. There is substantial support that APC may fuel the hyperactive immune response in human autoimmune disease by driving activation and differentiation of auto-reactive effector T-cell populations^14,15^. Recruitment of specific DC and/or monocytes in inflamed tissues may play a key role in the pathophysiology of the disease promoting T helper cell (Th) polarization depending on the environment. In JIA (the most common inflammatory arthropathy in children), severe chronic inflammation in the synovium leads to joint destruction and additional extra-articular manifestations (such as uveitis), as well as loss in quality of life for patients (pain and reduced functional ability in daily life)^16^. Sessions with joint puncture, aspiration and intra-articular corticosteroid injection are part of early clinical management of JIA, providing access to SF exudate which can be used for quantification and analysis^16^.

When compared to healthy controls, a study showed that patients with JIA have decreased PB DCs which are associated with a poor clinical outcome^17^. In contrast, an enrichment of cDCs is seen in the SF from inflamed joints, with a substantial expression of maturation markers, suggesting initiation and maintenance of inflammation^17^. As mentioned above, local environments and stimuli can initiate distinct modification to DC phenotype and function. When examining JIA patient SF exudate samples, a recent study found that total T cell, pDCs and cDC1s were increased with the latter being barely detectable in PB samples^7^. Similarly, in the inflamed synovial joint of rheumatoid arthritis (RA) patients, cDC1s were found to be enriched and capable of activating CD4^+^ and CD8^+^ T cells, contributing to inflammation^18^.

A working inflammatory arthropathy (IA) model (based on (pediatric and adult) human and murine models) suggests that autoantigen specific CD4+ T cells primed by DC enhances T cell maturation and presentation of neo-epitopes (auto-antigens). As such, autoreactive and potentially cross-reactive polarized T cells propagate autoimmune arthritis, contributing to the effector phase of the disease^19^. Interestingly, at the site of inflammation in chronic arthritis, the ability of SF-derived DC from RA patients to stimulate autologous PB T cells was shown to be superior to both PB-derived DC, and PB- and SF-derived monocytes^20^. Additionally, cDC2 from SF were demonstrated to be able to induce T cells to proliferate, and produce IFNγ and IL-17^21^, emphasizing their pro-inflammatory nature. Remarkably, in SF, there is a very large contribution of cDC1 to the pool of DC (∼35% compared to 3% in PB)^22^. cDC2 were also shown to be increased in SF (64% SF vs. 33% in PB), while pDC showed a decrease (33% PB vs. 9% SF)^22^. In steady state, cDC1 have been shown to be the counterpart of the murine CD8α^+^ and CD103^+^DC^23^. However, while murine CD8α^+^ DC have been demonstrated to be the most efficient at cross-presentation of all APC, human cDC1 have superior exogenous and necrotic antigen cross presentation and produce increased amounts of type III interferons^23–25^. Additional functions including IFNλ production, induction of cytotoxic T lymphocytes (CTL)^26^, and potential tolerogenic features^27^, suggest that cDC1 may play specific roles in immune responses.

The role of cDC1 has not been extensively described in reference to the pathogenesis of IA in general, but they were demonstrated to be distinct from PB counterparts and are capable of inducing CD4^+^ T cell responses and activating synovial fibroblasts^18^.

Here, we aimed to determine whether there is distinct functional programming of specific human APC subsets in synovial fluid from inflamed joints of JIA patients. We investigated their gene expression profile and performed extensive phenotyping and functional assays of the APC subsets to correlate the transcriptomic signatures to functional outcome. We show that while cDC2 and monocytes at the site of inflammation have a related but non-overlapping pro-inflammatory profile, the enriched cDC1 are more quiescent and less pro-inflammatory. Together the data demonstrate differential responsiveness and functional programming of APC subsets under inflammatory conditions.

## Materials & Methods

### Patients

32 patients with JIA according to the revised criteria for JIA^28^, were included in this study. Additionally, 4 RA patients were included. All patients had active disease and underwent therapeutic joint aspiration at the time of inclusion. 2 patients were later diagnosed as borrelia burgdorferi positive. Patients were between 2 and 18 years of age and were either untreated or treated with disease modifying anti-rheumatoid drugs (DMARDs), non-steroidal anti-inflammatory drugs (NSAIDs), methotrexate (MTX) or a combination (Supplementary Table 1). Additional clinical information can be found in Supplementary Table 1. Informed consent was received from parents/guardians and/or from participants directly when they were over 12 years of age. The study was approved by the Institutional Review Board of the University Medical Center Utrecht (UMCU) METC No.11-499/C (Pharmachild Study) and performed according to the principles expressed in the Helsinki Declaration.

### Definition of APC subsets

We defined APC as HLA-DR^+^, lymphocyte lineage-negative, and based on lineage-imprinted surface markers we delineated pDC (CD11c^−^ CD123^+^). Since monocytes can express many markers used to identify cDC, and therefore have often contaminated cDC populations, a strict separation based on CD14 and CD16 expression was applied. Here, CD14^+^ CD16^−/+^ cells were defined as monocytes (HLA-DR^+^CD11c^+^CD14^+^CD16^+^), while in the CD14/CD16-negative population we could define two types of cDC: cDC1 (HLA-DR^+^CD11c^+^CD14^−^CD16^−^ CD1c^−^CD141^hi^), and cDC2 (HLA-DR^+^CD11c^+^CD14^−^ CD16^−^ CD141^−/dim^CD1c^hi^) (Fig. 1A). Both cDC and monocytes expressed CD11c. Based on their relationship to each other, and the relatively big difference with pDC in terms of phenotype and described functions^2^, we decided to focus only on monocytes and cDC. Applying an unsupervised computational technique to visualize single-cell data and enable cellular hierarchy inference among subpopulations of similar cells (using SPADE^29^), the same APC populations were identified (Supplementary Figure 1A), supporting our sorting strategy.

**Figure 1.**
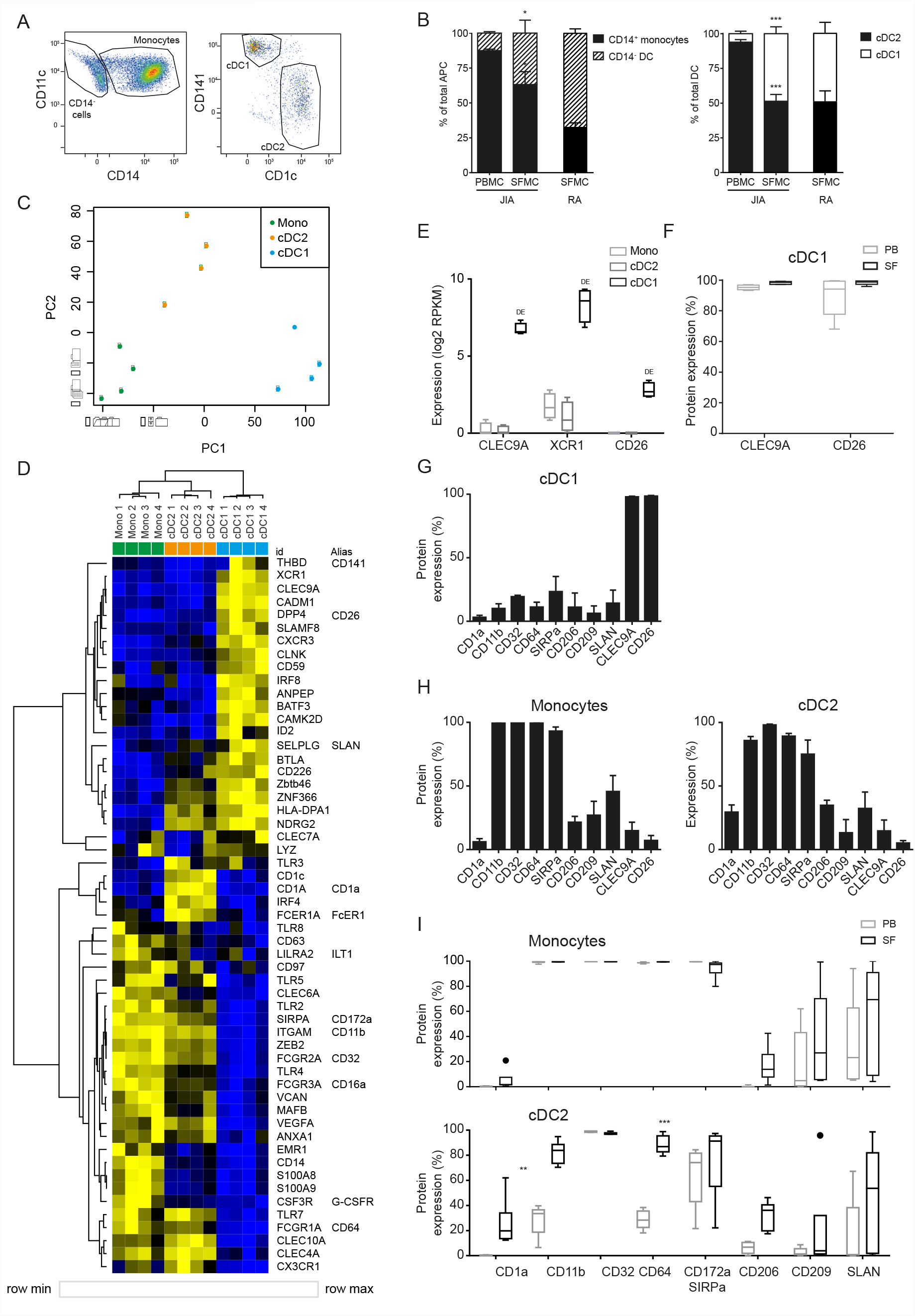
cDC1 are highly enriched in inflammation and distinct from other APC. (**A**) Example FACS plot showing the relative proportions of SF APC. HLA-DR^+^ cells were gated, and further subdivided into CD11c^+^ CD14^+^ monocytes, and CD11c^+^ CD14^−^ cDC, that were either type 1 or type 2. These cell types were quantified in (**B**) as monocyte/DC proportion within total APC (left panel), and cDC2/cDC1 proportions within total cDC (n=5). (**C**) PCA based on variable gene expression between SF APC subsets in RNAseq data. (**D**) Heatmap depicting relative gene expression of genes that were, based on literature, associated with monocyte-derived cells and cDC. Cluster analysis performed on both rows and columns (one minus Pearson correlation, average linkage) (**E**) Comparison of the gene expression of cDC1-related genes between SF APC subsets (n=4; DE, Differentially Expressed) (**F and I**) Expression of cDC1-related markers (F and I) and monocyte-related markers (I) measured by flow cytometry, compared between paired PB and SF samples (n=3 for CD32, n=5 for other markers; One-way ANOVA, ** p<0.01, ***p<0.001). (**G and H**) Expression of monocyte-related markers and cDC1-related markers, measured by flow cytometry (n=4 for CD32, n=9 for other markers).

### RNAseq

Total RNA was extracted using TRIzol reagent (Invitrogen) according to the manufacturer ‘s instructions and stored at −80°C until further processing. RNA yield was assessed by Qubit RNA BR assay kit and then integrity was determined by Bioanalyzer using the picochip. mRNA was selected using poly(A) purist MAG kit (Life Technologies) and additionally purified with a mRNA-ONLY™ Eukaryotic mRNA Isolation Kit (Epicentre). Transcriptome libraries were then constructed using SOLiD total RNA-seq kit (Applied Biosystems) and sequenced using 5500 W Series Genetic Analyzer (Applied Biosystems) to produce 40-bp-long reads.

### RNAseq data analysis

Sequencing reads were mapped against the reference human genome (hg19, NCBI37) using Burrows-Wheeler Aligner (BWA) (-c –l 25 –k 2 –n 10)^30^. Data were analyzed using DEseq2^31^ and custom Perl (www.perl.org) and R (www.r-project.org) scripts, including principal component analyses (PCA) (Fig. 1C, and Supplementary Figure 1F), unsupervised cluster analyses (Fig. 2E, Supplementary Figure 1A, and 1G), three-way comparisons in spider-plot format (Fig. 2B-D, and 3A), and k-means analyses (Fig. 4A, Supplementary Figure 4A, and 4C). Additional unsupervised cluster analyses (Fig. 1D, 4D, 4F, Supplementary Figures 1C-D, and 3A) were performed making use of Morpheus (https://software.broadinstitute.org/morpheus/). Using Morpheus ‘ standard settings, samples were clustered based on their Pearson distance, using average linkage. The Venn diagram in Fig. 2A was produced using jvenn (GenoToul, Toulouse)^32^, based on gene lists from two-way comparisons between SF monocytes and cDC subsets. Gene ontology based on gene lists derived from the abovementioned Venn diagram (for Fig. 2B-D, and 3A) and from k-means analysis (for Figure 4B, Supplementary Figure 4A-C), was determined using ToppFun, which is a part of the ToppGene Suite^33^.

**Figure 2.**
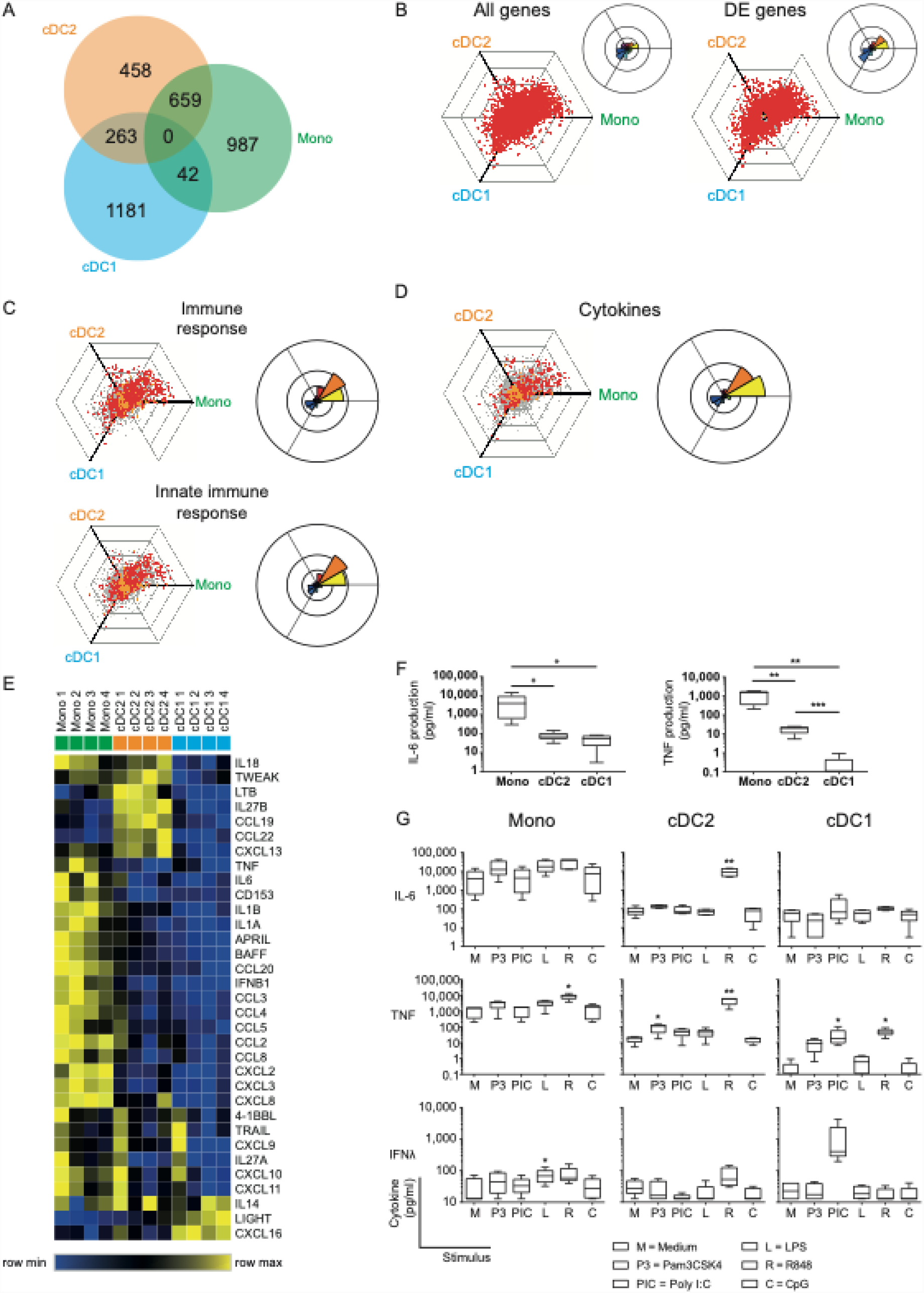
SF cDC2 and monocytes, and not cDC1, express an inflammatory phenotype. (**A**) Venn diagram depicting RNAseq DE genes as distributed over the SF APC subsets. (**B-D**) To visualize differential gene expression between SF APC, the expression of each gene by each cell type was vectorized and plotted in a hexagonal diagram, in which the direction of a point relative to the origin depicts an upregulation in the cell type(s) on that side of the hexagon, while the distance from the origin depicts the strength of this upregulation. Rose diagrams (right upper corner of or next to a hexagon) show the quantification of the genes in each direction. (**B**) Hexagon and rose diagrams of all genes measured (left panel), and all DE genes. (**C-D**) Hexagon and rose diagrams of DE genes measured (red dots) that, as defined by Gene Ontology annotation, have been associated with the immune response (**C**, top panel), the innate immune response (**C**, bottom panel), and cytokine production (**D**). Orange dots are associated non-DE genes, grey dots are unassociated non-DE genes. (**E**) Heatmap depicting relative gene expression of cytokine genes that were >1 log2 RPKM and DE between SF APC – cluster analysis performed on rows (one minus Pearson correlation, average linkage) (**F-G**) Cytokine production by *ex vivo* SF APC, as measured by Luminex (n=5). One-way ANOVA, * p<0.05, ** p<0.01, *** p<0.001. (**F**) Spontaneous production of IL-6 and TNF production by SF APC during overnight culture without further stimulation. (**G**) Spontaneous (M) or toll-like receptor (TLR) ligand-induced cytokine production by SF APC.

**Figure 3.**
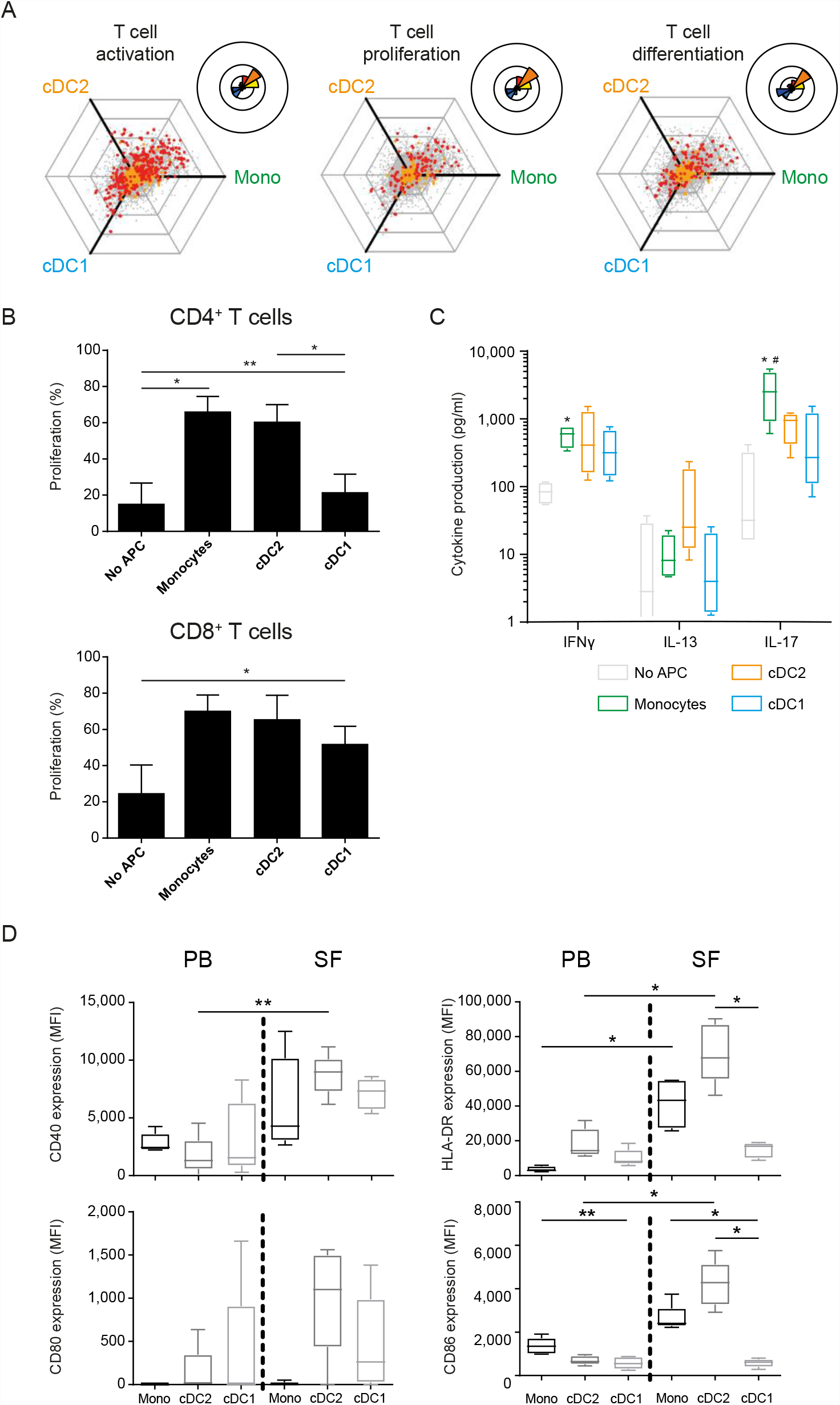
Functionally, cDC1 are weak inducers of T cells compared to cDC2 and monocytes. (**A**) Hexagon and rose diagrams of DE genes measured (red dots) that, as defined by Gene Ontology annotation, have been associated with T cell activation (left panel), T cell proliferation (middle panel), and T cell differentiation (right panel). **(B-C)** Proliferation of CD4^+^ (top panel) and CD8^+^ (bottom panel) T cells, as measured by flow cytometry (**B**, n=5), or cytokine production by CD3^+^ T cells, as measured by Luminex (**C**, n=4), after co-incubation of CD3^+^ cells with either no APC or one of three SF APC types for 5 days, in the presence of anti-CD3. (**D**) Expression of co-stimulatory markers CD80, CD86, MHC II molecule HLA-DR, and activation marker CD40 by APC in paired PB and SF, as measured by flow cytometry (n=5). One-way ANOVA, * p<0.05, ** p<0.01.

**Figure 4.**
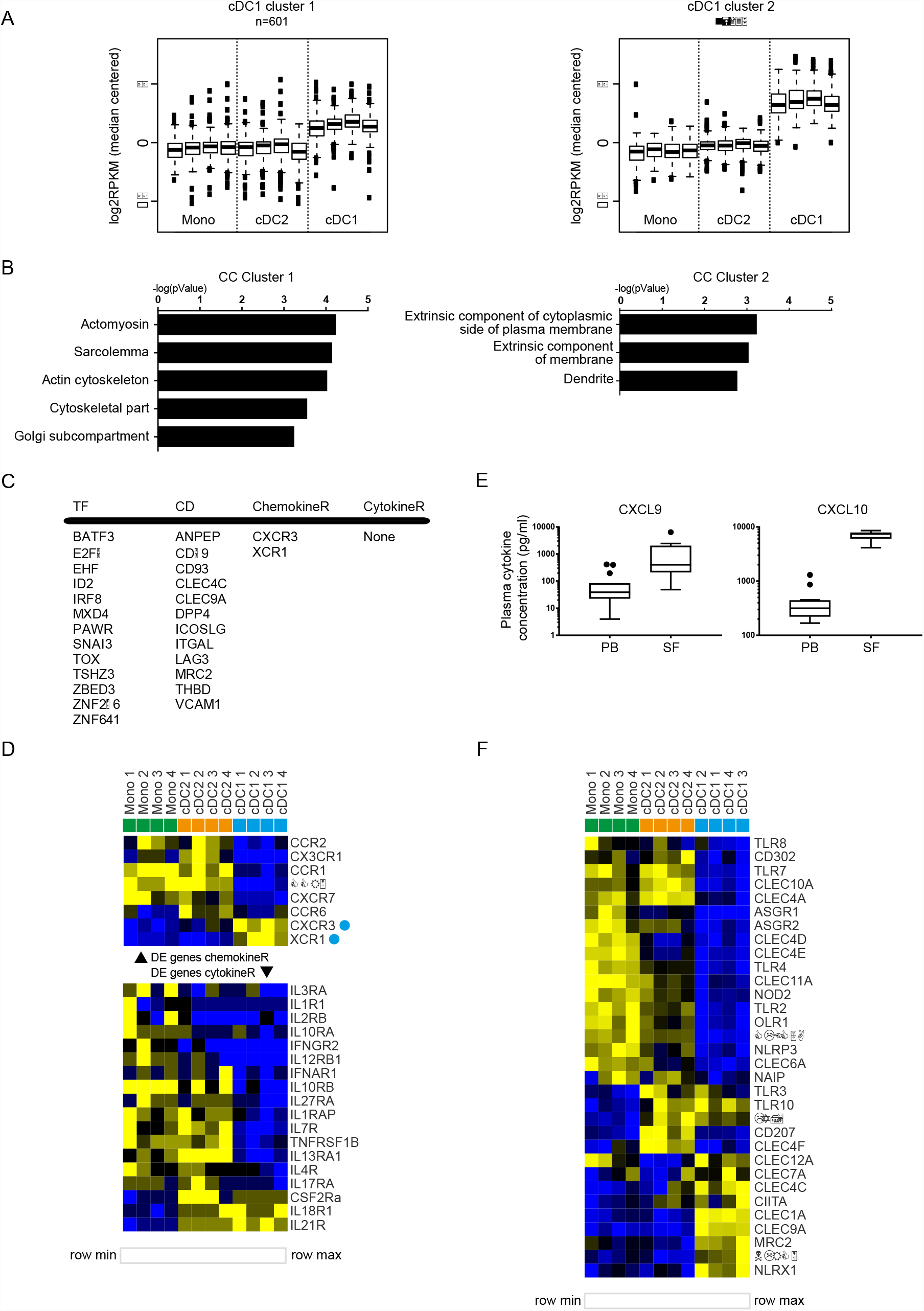
SF cDC1 present a quiescent phenotype, and a limited palette of receptors. (**A**) k-means clustering of SF APC RNAseq data, depicting clusters that contain genes highly (cluster 1) and very highly (cluster 2) expressed by cDC1 compared to cDC2 and monocytes. (**B**) GO terms associated with cluster 1 and cluster 2 genes, ranked by enrichment scores. (**C**) Selected DE genes high in cDC1. TF, transcription factors; CD, cluster of differentiation; ChemokineR, chemokine receptors; CytokineR, cytokine receptors. (**E**) Comparison of CXCL9 and CXCL10 levels between PB plasma samples and SF plasma samples (n=13-15). (**D & F**) Heatmaps depicting relative gene expression of chemokine receptor (E, top part), cytokine receptor (E, bottom part), and pattern recognition (F) genes that were >1 log2 RPKM and DE between SF APC (n=4) – cluster analysis performed on rows (E and F) and columns (F)(one minus Pearson correlation, average linkage).

For: *PBMC and SFMC isolation, Flow cytometric characterization and sort procedure of APC, APC TLR stimulation and cytokine measurement, Proliferation assays and Luminex*, please see the Supplementary Information section

### Statistics

Paired t-test (two-tailed) statistical tests were performed to assess possible differences between PB and SF APC percentages (Fig. 1B). One-way ANOVA statistical tests were used to assess differences in protein marker expression, cytokine production, the induction of T-cell proliferation and cytokine production, and plasma cytokine levels. When ANOVA testing yielded significant results, post-hoc tests were performed: Tukey ‘s post-hoc tests were performed for assessing differences in cytokine production between APC subsets (Fig. 2F) and between control and TLR-ligands (Fig. 2G), differences in T-cell proliferation induction between APC subsets (Fig. 3B), and differences in co-stimulatory molecule expression between APC subsets, and between PB and SF (Fig. 3D). Dunnett ‘s post-hoc tests were performed for assessing differences in protein marker expression between PB and SF (Fig. 1F, 1I and Suppl. Fig. 1E), differences in induction of T-cell cytokine production between control and APC subsets (Fig. 3C), and differences in cytokine levels between PB and SF plasma (Fig. 4D).

## Results

### Human cDC1 are highly enriched in SF and phenotypically distinct from other APC

To characterize human APC under inflammatory conditions, we analyzed APC subpopulations in SF obtained from inflamed joints of juvenile arthritis patients and compared these to paired peripheral blood samples. Within HLA-DR^+^CD11c^+^lineage^−^ cells we distinguished 3 subpopulations for further analysis: cDC1, cDC2, and CD14^+^ monocytes (Fig. 1A).

In SF the percentage of CD14^−^ DC from total APC was significantly higher than that in PB, although the majority still consisted of CD14^+^ monocytes (Fig. 1B, left panel). Furthermore, in SF we found a striking enrichment of cDC1 resulting in percentages comparable to those of cDC2 (Fig. 1B, right panel). Similar ratios of cDC subsets (Fig. 1B, right panel) were observed in SF-derived from inflamed joints of adult RA patients, indicating that our findings are not JIA-specific.

Next, we performed gene-expression profiling by RNA sequencing. Principal component analysis revealed that each APC subset plots separately, while the four replicates of each subset clustered together (Fig. 1C). In this PC1-PC2 2D space, monocytes and cDC2 were closer to each other, indicating higher similarity. Unsupervised clustering analysis also demonstrated that cDC1 are more distinct from the other two subsets, while cDC2 and monocytes, although clustering separately, share certain features (Supplementary Figure 1B). We then further compared these subsets based on published markers specific for DC subsets in steady state^5,10,11,13,34^. These markers could indeed separate the three APC subsets, based on hierarchical clustering (Fig. 1D). In concordance with the variable genes cluster analysis (Supplementary Figure 1B), cDC2 and monocytes partly overlapped in expression of certain genes (Fig. 1D). When directly comparing cDC2 and monocytes without cDC1 (Supplementary Figure 1C), it is well possible to separate these two cell types. Since inflammation could influence marker expression, we compared expression during steady state and inflammation: 4 out of 5 genes were specific for cDC2 in inflammation, and 5 out of 13 genes remained specific for monocytes during inflammation (Supplementary Figure 1D; underlined). Based on our data, a signature of nine markers can optimally distinguish between monocytes and cDC2 in both inflammation and steady state (cDC2: *IRF4, CD1c, FCER1A, CLEC10A*; monocytes: *ZEB2, FCGR2A, TLR4, MAFB, EMR1*). Additional phenotyping was performed by assessing protein expression of various surface markers by flow cytometry (as reviewed by Schlitzer et al.^3^). SF cDC1 expressed hallmark proteins CLEC9A, XCR1, and CD26 (Fig. 1E, mRNA and F, protein), while hardly expressing any of the surface markers strongly upregulated in SF monocytes and cDC2 (Fig. 1G-H). Neither the expression of CLEC9A and CD26 nor monocyte-related markers differed between PB and SF (Fig. 1F and Supplementary Figure 1E).

In contrast, the expression of monocyte-related markers was lower in PB cDC2 than in their SF counterparts (Fig. 1H, I). However, our principal component analysis and clustering analysis, both taking into account thousands of genes in addition to these APC markers, demonstrated that both PB and SF cDC2 are closer to each other than they are to PB monocytes (Supplementary Figure 1F-G).

### SF enriched cDC2 and monocytes display a clear pro-inflammatory phenotype, in contrast to SF enriched cDC1

Since the APC subsets were derived from a local inflammatory site, it is tempting to denominate these cells as inflammatory DC and inflammatory monocytes as has been done for other myeloid cells from inflammatory locations^21,35,36^. We compared the three SF subsets by performing pairwise comparisons, and a list of differentially expressed (DE) genes was comprised, with at least a 2-fold difference (P<0.05), between the different subsets (numbers of DE genes in Supplementary Table 2). All APC express comparable numbers of DE genes as shown by the Venn diagram (Fig. 2A), while DE genes specific for a subset or shared between subsets tend to differ per subset as displayed by the combined Venn diagram.

We next performed a three-way comparison of the gene expression profiles of the three cellular subsets, similar to the method described by Van de Laar et al^37^. In these hexagon diagrams the direction of each dot relative to the origin represents the vectorized expression of a gene in one or two subsets, where the distance from the origin represents the magnitude of the expression. Plotting all 32,584 unique transcripts shows that most genes are relatively equally expressed by the three subsets (Fig. 2B, left panel). When solely assessing the 3590 DE genes, vectorized expression is biased towards differences between cDC1 cells and either of the other two subtypes suggesting these are more dissimilar (Fig. 2B, right panel). When specifically analyzing genes associated with DC immune functions and innate immune responses, DE genes are mostly distributed over the cDC2/monocyte side of the diagrams (Fig. 2C-D).

In support of this, the pro-inflammatory cytokine and chemokine expression at the single gene level, including TNF, IL-6, IL-1β, CCL3, was highest in monocytes (Fig. 2E). Furthermore, the cytokine/chemokine signature clearly differed between the 3 subsets. Direct *ex vivo* culture of sorted subsets confirmed that the classical pro-inflammatory cytokines TNF and IL-6 were mostly produced and secreted by monocytes without any further stimulus needed, whereas cDC2 and especially cDC1 produced little of these cytokines spontaneously (Fig. 2F). However, the cDC2, were able to produce these and other cytokines (including IL-12 family members) upon specific TLR stimuli, especially Poly I:C and R848 (Fig. 2G, Supplementary Figure 2A).

Also, monocytes responded to these TLR ligands by further increasing their cytokine production (Fig. 2G, Supplementary Fig. 2A). cDC1 solely reacted to Poly I:C and R848 that induced low levels of TNF-α production. To show that cDC1 are still functional, upon Poly I:C stimulation, the cDC1 selectively produced high levels of IFN λ (Fig. 2G), one of the hallmark cytokines that have been attributed to cDC1^38^, demonstrating functional differences between these APC subsets.

Taken together, these data demonstrate both phenotypical and functional differences of the specific APC subsets at the site of inflammation when compared to peripheral blood. Here, cDC1 lack the pro-inflammatory phenotype and function displayed by the cDC2 and monocytes that were derived from the same inflammatory environment.

### cDC1 are weak inducers of T cell proliferation and activation compared to cDC2 and monocytes

To further dissect functions of APC subsets in inflammation we assessed their capacity to regulate T cell function. In addition to their pro-inflammatory status, cDC2 and monocytes demonstrated higher expression of genes associated with T cell activation, proliferation and differentiation, compared to cDC1 (Fig. 3A). To confirm their ability to induce T cell activation, we incubated them with autologous PB-derived T cells. Indeed, in contrast to cDC2 and monocytes, cDC1 were weak inducers of T cell proliferation, especially of CD4^+^ T cells (Fig. 3B). All subsets equally induced the production of the T_H_1-associated and CD8^+^ T cell-associated cytokine IFNγ, whereas the T_H_2 cytokine IL-13 was predominantly induced by cDC2 (Fig. 3C). Only monocytes strongly induced IL-17, the hallmark cytokine of T_H_17 cells and linked to pathogenesis and damage in autoimmune arthritis^39^.

Analyzing all co-stimulatory, co-inhibitory, and dual-activity molecules that were DE between the subsets (Supplementary Figure 3A) we found that most of these molecules were preferentially expressed in monocytes and cDC2. This was confirmed for a selection of molecules on the protein level (Fig. 3D). Altogether, in contrast to SF cDC1, SF cDC2 and monocytes are strong inducers of T cells, likely as a result of high expression of cytokines and co-stimulatory proteins.

### The quiescent phenotype of SF cDC1 is associated with a limited palette of pattern recognition receptors (PRR), cytokine and chemokine receptors

To further assess the cDC1 at the site of, we performed k-means clustering and selected those clusters with high gene expression selectively in cDC1 (Fig. 4A). While genes expressed highly in cDC2 and monocytes were associated with immune-related functions (Fig. 2C-D, 3A and Supplementary Fig. 4A), cDC1-related genes (of both clusters) did not yield many annotations nor did they associate with immune functions (Fig. 4B and Supplementary Figure 4B). For instance, cDC1 enriched genes more often associated with cytoskeletal functions and the plasma membrane (Fig. 4B). Interestingly, one of the clusters presented a cDC signature, containing genes that were highly expressed in both cDC subsets compared to monocytes (Supplementary Figure 2C). The genes in this cluster associated with immune response functions, especially antigen-presentation and co-stimulation, albeit to a much lesser degree than similar functions associated with cDC2/monocyte high genes (Supplementary Figure 4A).

Assessing the cDC2-high gene list, Figure 4C depicts a selection of upregulated genes. This includes factors with a well-defined role in cDC1 development, such as BATF3, ID2 and IRF8. Likewise, 4 of 12 CD molecules have been described as cDC1-specific: THBD (CD141), DPP4 (CD26), ANPEP (CD13), CLEC9A (CD370). To find clues for the factors underlying the preferential enrichment of cDC1 in SF, we assessed molecules involved in migration of immune cells. Only two chemokine receptors were specifically highly expressed by cDC1 namely CXCR3 and XCR1 (Fig. 4C,D). Interestingly, analyzing plasma concentrations for several ligands we found that the levels of two of the ligands for CXCR3, CXCL9 and CXCL10, were highly increased in SF compared to levels in PB (Fig. 4E). These data suggests that not the subset-specific XCR1, but instead a more broadly expressed receptor like CXCR3 may contribute to the accumulation of cDC1 in SF.

Additionally, the expression of pattern recognition receptors, especially TLR, was also lower in cDC1 compared to cDC2 and monocytes (Fig. 4F), but also low in expression in an absolute sense (data not shown), together indicating a relatively specified potential to respond to the local stimuli. Altogether these data show a relative quiescence of cDC1 in SF compared to cDC2.

## Discussion

Until recently, monocytes and DC have mostly been characterized in steady state, however it remains unclear how these cells – especially in humans – are programmed under inflammatory conditions^2,7,10,13,17,18^. To fill this gap in knowledge, we have comprehensively profiled monocytes and cDC subsets in the exudate derived from the inflamed joint of JIA patients. Although SF from the joints of arthritis patients, may not be a complete reflection of the inflammation taking place in the tissue, it has proven to be an excellent model to study APC in a highly inflammatory environment^7,21^. Our findings show that at the site of inflammation, there is specific, functional programming of human DC subsets. In this context both monocytes and cDC2 demonstrate a clear and distinct pro-inflammatory role whereas the enriched cDC1 remain surprisingly quiescent on both a transcriptional and functional level. Despite some overlap in transcriptional profile and phenotype, we propose that inflammatory cDC2 are not monocyte-derived, but *bona fide* conventional DC. Based on principal component analysis and unsupervised clustering analysis, CD14, CD16, CD1c expression, but also IRF4 and IRF8 levels can be markers used to further distinguish cDC and monocytes^11^. Recently, additional classifications have been made to DC subsets. A study by Villani et al^10^ demonstrated that, in steady-state, out of six newly described DC subsets, sub-divisions of the CD1C/BDCA-1^+^ cDC2 delineate DC2 (CD5^+/-^) and DC3 (cD1c^+^CD5^−^CD163^+^CD14^lo-hi^), although we have not yet incorporated these distinctions into our research.

In our transcriptional data-set, both SF cDC2 and SF monocytes were very similar to their PB counterparts, indicating these SF subsets likely originate from the blood and did not differentiate from other APC subsets (monocytes) at the site of inflammation. This may seem in contrast to what Segura et al.^20^, described earlier, showing a population of inflammatory DC in RA SF and inflammatory tumor ascites that they proposed to be monocyte-derived^20^. This distinction may be explained by a difference in DC sorting strategy. We indeed observed CD1c expression on SF monocytes in our JIA patients, albeit at lower levels than on the CD14^−^CD1c^+^ selected population and this was confirmed on the mRNA level. Supporting this hypothesis, our SF CD14^+^ monocyte population, similar to the inflammatory DC described in the paper by Segura et al^20^, produced the highest levels of the T_H_17-promoting cytokines IL-6, TNF, IL-23, and IL-1β and demonstrated the strongest induction of IL-17 production by T cells. In addition, these cells were distinguished by strong expression of amongst others acute and chronic inflammatory genes, and had strong T cell-activating capacity^10^. In contrast, CD14^+^ CD1c^+^ DC have been shown in blood of healthy controls and melanoma patients, albeit showing immune-inhibitory functions^40^. In cancer settings, it was also demonstrated that pre-cDC1, lineage-committed progenitors that give rise to cDC1, express the T helper type 1-associated chemokine receptor CXCR3 for homing functions to the melanoma. Preservation of CXCR3-ligands CXCL9 and CXCL10 promoted cDC1 presence in tumors, highlighting a mechanism that may reflect similarly increased levels of CXCL9 and CXCL10 in SF^41^.

The expression profile of cDC1-associated markers on a transcriptional and protein level confirmed that our SF-derived CD141^+^ cells consisted of *bona fide* cDC1^38,42,43^. This was further supported by the limited response to a number of TLR ligands and specifically, the production of IFNλ upon poly I:C exposure – cDC1 features that both have been described before in steady state^37,44^. Together, our findings suggest that although the APC subsets undergo specific inflammation-induced programming and differentiation/activation, the different APC subsets are still clearly distinguishable based on their steady state-defined core signature. Single-cell sequencing will shed further light on the heterogeneity and differentiation within the subsets.

We here demonstrate that SF cDC1 did not show a strong pro-inflammatory profile and induced only weak T cell proliferation and cytokine production compared to the other DC subsets One reason for this relative unresponsiveness may be the low expression of PRR, chemokine, and cytokine receptors on the SF CD141^+^ cDC that we observed. cDC1 have been reported to play a major role in cross-presentation, i.e., the presentation of acquired exogenous antigens via MHC class I which is essential for the initiation of CD8+ T cell responses^24,45^. The low MHC class I expression by cDC1 in our study may affect CD8 T cell responses, but that should be assessed with antigen-specific T cell responses and preferably also in a cross-presenting setting. However, although cross-presentation is one of the key features of cDC1 in mice, in humans it is not restricted to the cDC1 lineage^21,23^. In mice, tolerance induction has been a proposed function of cDC1, by inducing Treg^25,46^, for example via the production of retinoic acid^25^. In humans, this potential has also been reported^27^, describing CD141+ DC that induced T cell hypo-responsiveness and potent regulatory T cells. It is not clear however whether these are *bona fide* cDC1 or other cells expressing CD141, since expression of other cDC1 markers like XCR1 was not shown, and the cells were also CD14^+^.

Additionally, these cells were found to produce IL-10 spontaneously, while the SF cDC1 in our study did not produce IL-10, in contrast to cDC2 and especially monocytes (data not shown). Furthermore, the aldehyde dehydrogenase genes 1-3 for the proteins needed for retinoic acid production^46^ were expressed by SF cDC1, but lower than or equal to that of the other subsets (data not shown), indicating that this may not be a major mechanisms of tolerance induction by SF cDC1. Interestingly, one cytokine that was restricted to the SF cDC1 was IFNλ. IFNλ was produced by SF cDC1 upon poly I:C stimulation, but was also found in SF plasma, as measured by Luminex, about 10x as high than that in PB plasma of patients with active disease (478.1±394.7 vs. 47.88±79.98, respectively; data not shown). This cytokine has been shown to induce the expansion of Treg^47,48^. Mennechet et al., showed that the effect was indirect via inducing tolerogenic monocyte-derived DC, that in turn induced proliferation of Treg^48^. Additionally, IFNλ has been suggested to have a dynamic in the progression of inflammation and autoimmunity specifically^49–51^.

Research to the implications of APC differentiation and specialization in the pathogenesis and progression of autoimmune diseases, especially inflammatory arthritis is gaining momentum^10,16,18,47^. Our data contribute to the understanding of functional contribution and dynamics of APC subsets in inflammatory environments. Both cDC2 and monocytes seem instrumental in the pro-inflammatory process whereas the relative quiescence of cDC1 in combination with their selective IFNλ production suggest a potential regulatory role for this enriched DC subset in tissue inflammation. Together, insights in the tolerance capacity of DC may be harnessed for potential immunomodulatory therapeutics in IA patients^52^, allowing for long-term, efficient treatment that can be tailored to patients ‘ unique cases.

## Author Contributions

Conceptualization: A.B., M.P., S.N., and F.W.; Selection of patients, acquisition patient data, synovial fluids and peripheral blood: J.F.S and B.J.V.; Methodology: A.B., and F.W.; Software: M.M.; Formal Analysis: A.B., M.M., B.C., and M.C.; Investigation: A.B.; Resources: A.B., A.S., and F.W.; Writing – Original Draft: A.B., A.S., and M.P.; Writing – Review & Editing: A.B., A.S., M.P., M.M., B.C., S.N., J.F.S., M.C., B.J.V., J.V.L., and F.W.; Visualization: A.B., A.S., M.P., and M.M., and B.C.; Supervision: F.W.; Funding Acquisition: F.W and J.V.L.

## Supplementary Figures

**Supplementary Figure 1.**
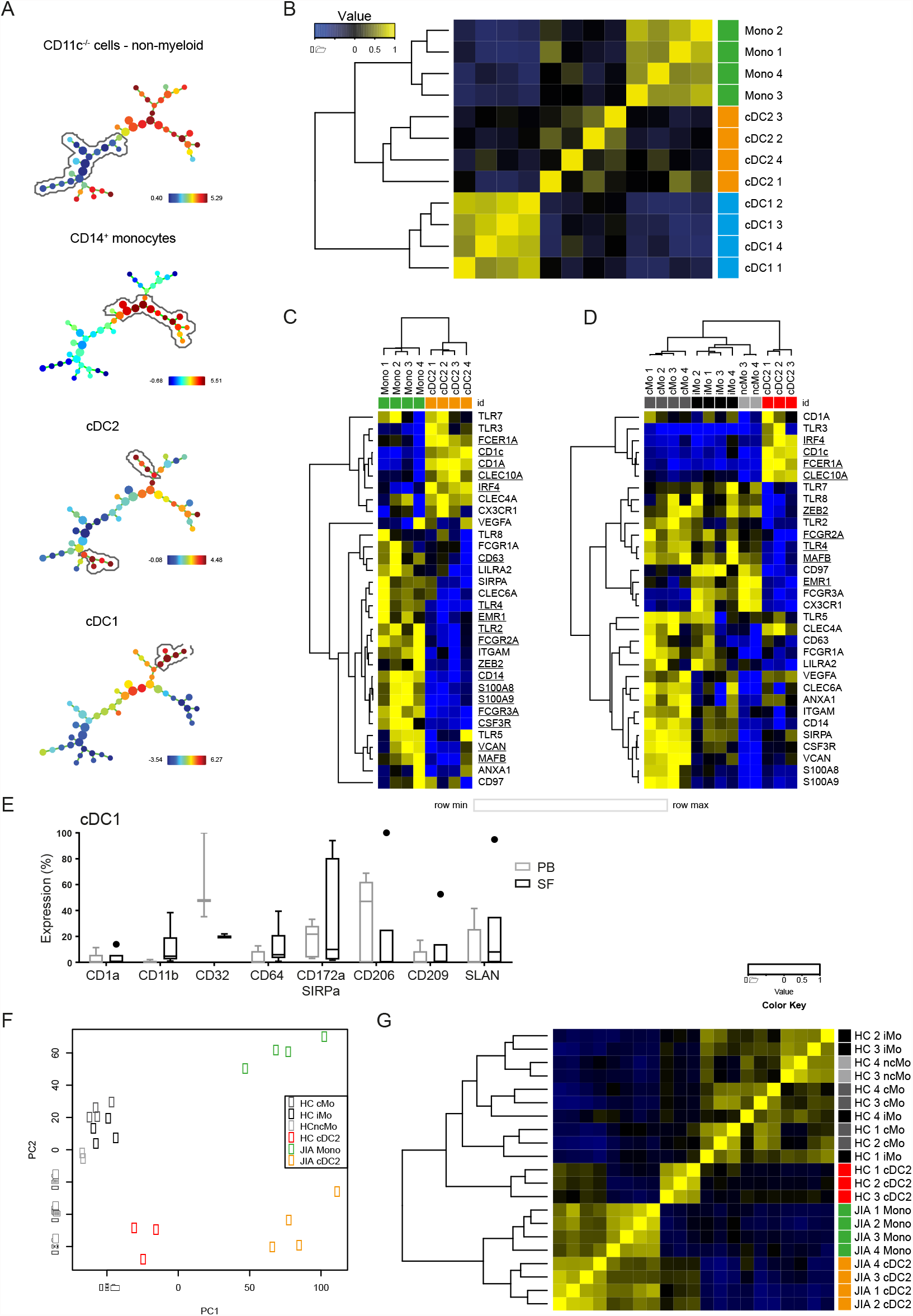
Related to Figure 1. (***A****) Unsupervised computational identification of APC populations using SPADE. Expression of the marker mentioned above each subfigure is quantitatively provided by means of color, while each subset mentioned is circled in its corresponding subfigure. (****B****) Unsupervised clustering heatmap of SF APC subsets. (****C****-****D****) Heatmaps* depicting relative gene expression of genes that were, based on literature, associated with monocyte-derived cells and cDC (CD141^+^ cDC-aligned markers from Fig. 1D removed) – cluster analysis performed on both rows and columns (one minus Pearson correlation, average linkage); (**C**) Comparison of SF monocytes and CD1c^+^ cDC; (**D**) Comparison of PB monocytes and CD1c^+^ cDC. cMo, classical monocyte; iMo, intermediate monocyte; ncMo, non-classical monocyte. (**E**) Comparison of monocyte-related markers and CD141^+^ cDC-related markers between paired PB and SF CD141^+^ cDC, as measured by flow cytometry (n=3 for CD32, n=5 for other markers). (**F**-**G**) PCA (**F**) and hierarchical clustering heatmap (**G**) based on variable gene expression between SF and PB monocytes and CD1c^+^ cDC in RNAseq data.

**Supplementary Figure 2.**
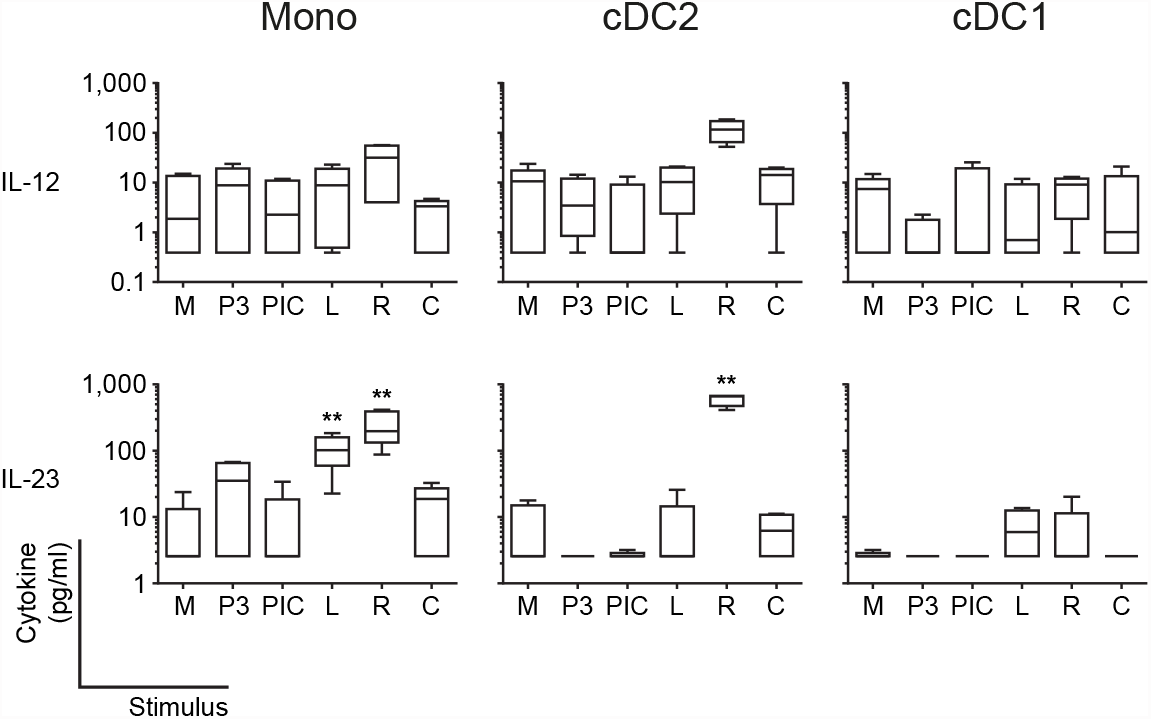
Related to Figure 2. (**A**) Spontaneous (M) or toll-like receptor (TLR) ligand-induced cytokine production by SF APC during overnight culture.

**Supplementary Figure 3.**
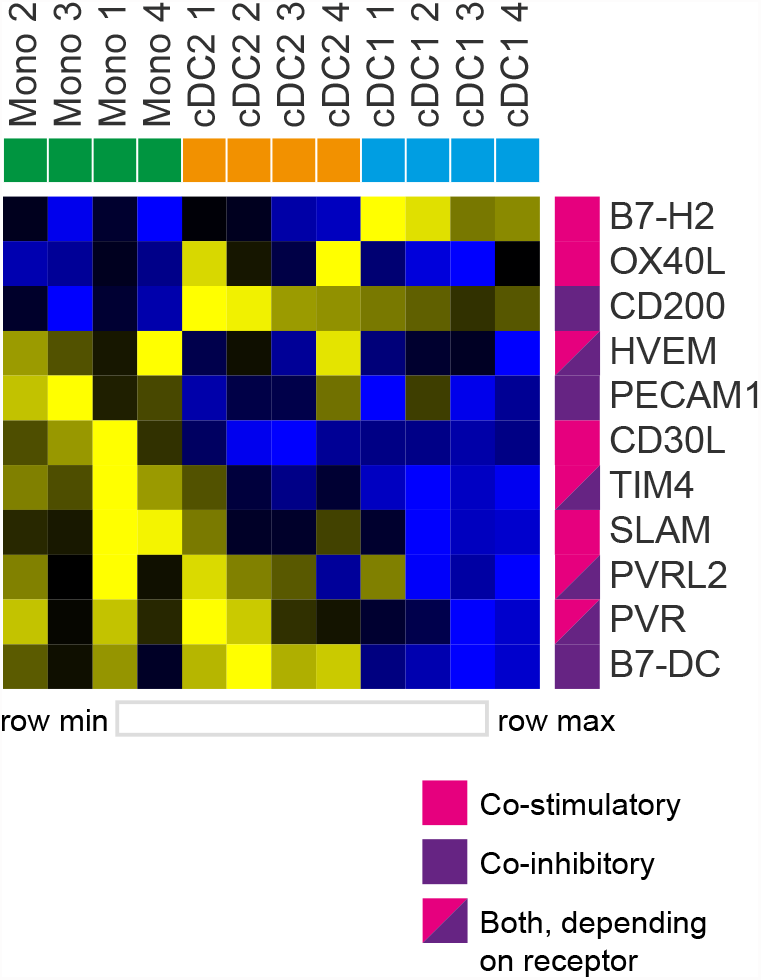
Related to Figure 3. (**A**) Heatmap depicting relative gene expression of co-stimulatory/-inhibitory genes that were >1 log2 RPKM and DE between SF APC – cluster analysis performed on both rows and columns (one minus Pearson correlation, average linkage).

**Supplementary Figure 4.**
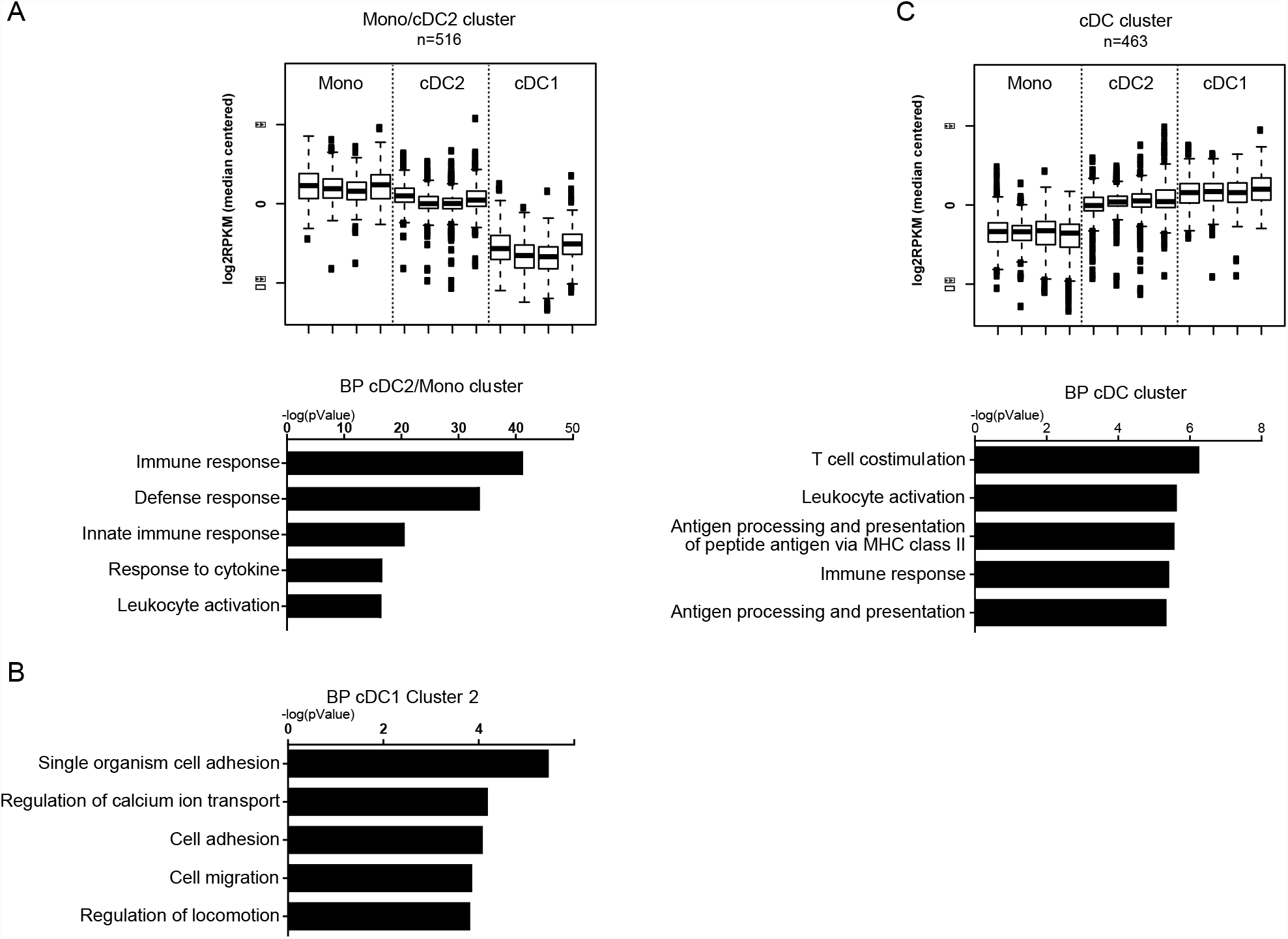
Related to Figure 4. (**A**) k-means clustering of SF APC RNAseq data (top panel), depicting a cluster that contain genes highly expressed by CD1c^+^ cDC and monocytes compared to CD141^+^ cDC, and GO Biological Processes (BP) terms associated with this cluster (lower panel), ranked by enrichment scores. (**B**) GO BP terms associated with CD141^+^ cDC cluster 2 genes, ranked by enrichment scores. (**C**) k-means clustering of SF APC RNAseq data (top panel), depicting a cluster that contain genes highly expressed by cDC compared to monocytes, i.e. a cDC cluster, and GO BP terms associated with this cluster (lower panel), ranked by enrichment scores.

**Supplementary Table 1:** Extended patient data including age, gender, type of JIA (abbreviations are: oligo-oligoarticular, poly-polyarticular, PsA-psoriatic arthritis), treatment status (abbreviations are: MTX-methotrexate, NSAID-non-steroidal anti-inflammatory drug) and (c)JADAS score – a physician-assigned, clinical score measuring overall disease activity.

| **Patient Index** | **Age (years)** | **Gender** | **Type of JIA** | **Treatment** | **(c)JADAS Score** |
| --- | --- | --- | --- | --- | --- |
| 1 | 9 | F | Enthesitis related-JIA | MTX | 8 |
| 2 | 16 | M | Oligo | None | 7 |
| 3 | 8 | M | Oligo | MTX | 7 |
| 4 | 12 | F | Extended oligo | None | 6 |
| 5 | 19 | F | Extended oligo | Adalimumab | 6 |
| 6 | 14 | M | Extended oligo | MTX | 4 |
| 7 | 11 | F | Oligo | MTX | 4 |
| 8 | 9 | F | Oligo | MTX | 1 |
| 9 | 10 | F | Poly | None | 4 |
| 10 | 15 | M | Oligo | Unknown | N/A |
| 11 | 14 | M | Oligo | None | 3 |
| 12 | 17 | F | Oligo | None | 5 |
| 13 | 16 | M | Oligo persistent | None | 3 |
| 14 | 9 | M | Poly JIA/PsA | MTX, Adalimumab | 6 |
| 15 | 15 | M | Oligo persistent | NSAID | 9 |
| 16 | 7 | F | Oligo, Uveitis | MTX | 3 |
| 17 | 8 | F | Oligo | Leflunomide | 2 |
| 18 | 14 | F | Undetermined JIA | NSAID | 6 |
| 19 | 12 | M | Oligo | None | 4 |
| 20 | 3 | F | Oligo | None | 7 |
| 21 | 7 | F | Oligo | None | 3 |
| 22 | 8 | M | Oligo | None | 5 |
| 23 | 5 | M | Oligo | None | 4 |
| 24 | 15 | F | Oligo | None | 4 |
| 25 | 9 | F | Oligo | None | 4 |
| 26 | 11 | M | JIA, not specified further | MTX | 3 |
| 27 | 8 | M | Oligo | None | 3 |
| 28 | 11 | F | Oligo | None | 8 |
| 29 | 17 | M | Oligo | MTX | 4 |
| 30 | 4 | F | Oligo | None | 4 |
| 31 | 12 | M | Oligo | None | 3 |
| 32 | 17 | F | Oligo | None | 5 |

**Supplementary Table 2:** 1807 DE genes between SF CD1c^+^ cDC and CD141^+^ cDC, of which 780 were higher in CD141^+^ cDC, while 1027 were higher in CD1c+ cDC. There were 1260 DE genes between CD1c+ cDC and iMo, 545 and 715 of which were higher expressed in CD1c+ cDC or iMo respectively. CD141+ cDC and iMo had a higher expression of 1283 and 1577 genes, respectively, resulting in 2860 DE genes.

|  | CD1c vs. CD141 | CD1c vs. Mono | CD141 vs. Mono |
| --- | --- | --- | --- |
| **CD1c high** | 1027 | 545 | - |
| **CD141 high** | 780 | - | 1283 |
| **Mono high** | - | 715 | 1577 |
| **Total DE** | 1807 | 1260 | 2860 |

## Supplementary Information – Methods & Materials

### PBMC and SFMC isolation

Peripheral blood (PB) was drawn via veni puncture. Synovial fluid (SF) was collected during therapeutic joint aspiration. Paired PB and SF samples were taken from the same patient at the same time. SF was first treated with hyaluronidase for 30min (37°C, 5% CO_2_) to reduce the viscosity, followed by centrifugation (4°C, 1600rpm).

Supernatants were removed for an additional centrifugation step (4°C, 3000rpm) to acquire SF plasma, while cell pellets were used for further SF mononuclear cells (SFMC) isolation. PB was first centrifuged (37°C, 1200rpm) to acquire PB plasma, while cell pellets were subsequently used for PB mononuclear cells (PBMC) isolation. Both SF and PB plasma were frozen at −80°C until later use. In order to obtain SFMC and PBMC, cell pellets were resuspended in RPMI containing 1% penicillin/streptomycin and cells were isolated using Ficoll-Paque density gradient centrifugation (GE Healthcare Bio-Sciences, AB) and were used either directly, or frozen in RPMI medium containing 20% FCS (Invitrogen) and 10% DMSO (Sigma-Aldrich) until further experimentation.

### Flow cytometric characterization and sort procedure of APC

To characterize the distribution and phenotype of APC subsets in PB and SF, fresh PBMC and SFMC were stained with antibodies against HLA-DR (Clone: G46-6; BD Pharmingen or L243; Biolegend), CD11c (3.9; eBioscience or B-ly6; BD Biosciences), CD123 (7G3, BD Biosciences), CD14 (M5E2; BD Biosciences), CD1c (AD5-8E7; Miltenyi or L161; Biolegend), and CD141 (M80; Biolegend or AD5-14H12; Miltenyi) to discern the respective APC subsets: pDC (HLA-DR^+^, CD11c^−/lo^, CD123^+^; not shown), monocytes (HLA-DR+ CD11c^+^ CD14^+^), CD1c^+^ cDC (HLA-DR^+^ CD11c^+^, CD14^−^, CD1c^+^ CD141^−/lo^), and CD141^+^ cDC (HLA^−^DR^+^ CD11c^+^, CD14^−^, CD141^+^ CD1c^−^), as shown in Figure 1A. Additional cellular hierarchy analysis from this cytometry data was performed using SPADE (Figure S1A), as described by its developers^1^. Specific settings used: FSC, SSC, CD141, CD1c, CD11c, CD14 as overlapping markers used for SPADE tree; apply compensation matrix in FCS header; Arcsinh with cofactor 150; number of desired clusters: 50.

Further phenotyping of subset markers (as shown in Figure 1E-F, 1H-I, S1G) and co-stimulatory molecules, and MHC molecules (Figure 3E) of each subset was performed using antibodies against CD1a (HI149; BD Biosciences), CD11b (ICRF44; eBioscience), CD32 (FLI8.26; BD Pharmingen), CD64 (10.1; Biolegend), CD172a (SE5A5; Biolegend), CD206 (19.2; BD Biosciences), CD209 (DCN46; BD Biosciences), SLAN (DD-1; Miltenyi), CLEC9A (8F9; Miltenyi), CD26 (2A6; eBioscience), CD40 (5C3, eBioscience), CD80 (L307.4, BD Biosciences), CD86 (IT2.2), and HLA-DR (L243, both Biolegend) and subsequently acquiring on FACSCanto II (BD Biosciences). Analysis was done using FlowJo version 10 (FlowJo, LLC).

To obtain APC subsets for functional assays or sequencing, above subset staining and gating was used to sort APC subsets on a FACSAria II or III (BD). In case of sorting APC subsets for RNAseq purposes, cells were additionally stained with antibodies against CD3 (OKT3; Biolegend), CD19 (HIB19; Biolegend), CD56 (HCD56; Biolegend) to gate out lymphocyte lineages before applying abovementioned gating strategy. After sorting, cells were washed in MACS buffer. If used directly for functional experiments, cells were resuspended in culture medium (RPMI 1640 (Lonza), 10% FCS). If intended for RNAseq purposes, sorted cells were dissolved in TRIzol (Invitrogen) and frozen at −80°C until further use.

### APC TLR stimulation and cytokine measurement

To measure cytokine production by SF-derived monocytes and dendritic cells subsets (Figure 2F-G, and Supplementary Figure 2A), APC subsets were sorted by flow cytometry as described above and 10.000 cells were then cultured in 100µl culture volume. Cells were either not stimulated or stimulated with Pam3CSK4 (100ng/ml), Poly(I:C) (25µg/ml), LPS (100ng/ml), CpG-A (5µg/ml; all Invivogen), or R848 (1µg/ml; Enzo). After overnight culture supernatants were collected and stored at −80°C until analysis.

### Proliferation assays

To assess the T-cell proliferation- and cytokine production-inducing capacity of APC subsets (Figure 3B-C), patient PB and SF samples were split into two fractions using CD3 magnetic-activated cell sorting (MACS) microbeads (Miltenyi Biotec), yielding CD3^+^ T cells, and CD3^−^ cells. CD3^−^ cells were subsequently stained for sorting APC subsets as described above. PB CD3^+^ T cells were labeled with 2µM cell tracer (Invitrogen) for 7 minutes at 37°C and extensively washed before used in proliferation assays. 50.000 CD3^+^ T cells were co-cultured with 10.000 SF-derived monocytes or DC subsets. At day 5, supernatants were collected to measure cytokine production, and cells were harvested, washed, and stained for T cell markers CD3 (Clone: SK7; BD), CD4 (RPA-T4; Biolegend) and CD8 (SK1; BD Biosciences). Proliferation of T cells (Figure 3B) was acquired by flow cytometry on a FACSCanto II (BD) and analyzed using FlowJo version 10 (FlowJo, LLC).

To measure APC-induced T-cell cytokine production (Figure 3C), supernatants were collected from proliferation assays and stored at −80°C until analysis.

### Luminex

Cytokine concentrations in PB plasma, SF plasma (both in Figure 4D), APC O/N stimulation supernatants (Figure 2F-G, and Supplementary Figure 2A) and APC-T cell co-culture supernatants (Figure 3C) were measured by Luminex technology as previously described^2^.

